# A cytokine network balance influences the fate of *Leishmania (Viannia) braziliensis* infection in a cutaneous leishmaniasis hamster model

**DOI:** 10.1101/2020.12.14.422666

**Authors:** MB Paiva, RP Ribeiro-Romão, L Resende-Vieira, T Braga-Gomes, MP Oliveira, AF Saavedra, L Silva-Couto, HG Albuquerque, OC Moreira, EF Pinto, AM Da-Cruz, A Gomes-Silva

**Affiliations:** Laboratório Interdisciplinar de Pesquisas Médicas, Instituto Oswaldo Cruz, FIOCRUZ/RJ; Laboratório de Biologia Molecular e Doenças Endêmicas, Instituto Oswaldo Cruz, FIOCRUZ/RJ; Laboratório de Pesquisa Clínica em Micobacterioses, Instituto Nacional de Infectologia Evandro Chagas, FIOCRUZ/RJ; Disciplina de Parasitologia-DMIP, Faculdade de Ciências Médicas, UERJ, Rio de Janeiro, Brazil; Laboratório de Doenças Parasitárias, Instituto Oswaldo Cruz, FIOCRUZ/RJ; Rede de Pesquisas em Saúde do Estado do Rio de Janeiro/FAPERJ; The National Institute of Science and Technology on Neuroimmunomodulation (INCT-NIM)

**Keywords:** Hamster, *Leishmania (Viannia) braziliensis*, cytokines, cutaneous lesions, parasite load, iNOS/arginase, immune-regulation, disease control

## Abstract

The golden hamster is a suitable model for studying cutaneous leishmaniasis (CL) due to *Leishmania (Viannia) braziliensis*. Immunopathological mechanismsare wellstablished inthe *L. (L.) major-mouse* model, in which IL-4 instructs a Th2 response towards progressive infection. In the present study, we evaluatedthe natural history of *L. braziliensis* infection from its first stagesup to lesion establishment, with the aim ofidentifyingimmunological parameters associated with the disease outcome and parasitismfate. To this end, hamsters infected with 10^4^, 10^5^,or 10^6^ promastigoteswere monitored duringthe first hours (4h, 24h), early (15, 30 days) and late (50 days) post-infection (pi) phases. Cytokines, iNOS and arginasegene expression were quantified in the established lesions by RT-PCR. Compared to the 10^5^ or 10^6^ groups, 10^4^animals presented lower lesions sizes, less tissue damage,and lower IgG levels. Basal gene expression in normal skin was high for TGF-β, and intermediary for TNF, IL-6, and IL-4.At 4hpi, no cytokine induction was observed in the 10^4^ group, while an upregulation of IL-6, IL-10, and IL-4 was observed in the 10^6^ group. At 15dpi, lesion appearance was accompanied byan increasedexpression of all assessed cytokines, markedly in the 10^5^ and 10^6^ groups. Upregulation of all investigated cytokines was observed in the late phase, although less expressive in the 10^4^ group. IFN-γ was the depending variable influencing tissue damage, while IL-6 was associatedto parasite load. The network correlating gene expression and clinical and laboratorial parameters indicated inoculum-independent associations at 15 and 30dpi.A strong positive network correlation was observed in the 10^4^ group, but not in the10^5^ or 10^6^ groups. In conclusion, IL-4, IL-6, IL-10, and TGF-β are linkedto *L. braziliensisprogression*. However, a balanced cytokine network is the key for an immune response able to reduce the ongoing infection and reduce pathological damage.

## Introduction

Tegumentary leishmaniasis (TL) is an infectious disease associated with poverty, and a general increasing trend in the number of new cases isreported annually to World Health Organization (1). The disease is caused by a protozoan belonging to the*Leishmania* genus, which compromises the skin and mucosa. Cutaneous leishmaniasis (CL) is the most common clinical form, characterized by one or more skin ulcers well delimited by a raised border at the site of the sand-fly infected bite. The *Leishmania* species implicated in CL can vary according to geographic region, and eightspecies havebeenidentified in Brazil (2, 3). Although most patients develop cutaneous ulcers, increasing evidence indicatesclinical presentation characteristics associated with a particular *Leishmania* species (4, 5, 6, 7). Because of this, the understanding of CL physiopathology mustbe particular to each *Leishmania* species. *L. (Viannia) braziliensis* is the most spread parasite in Brazil, causing not only CL but also mucosal leishmaniasis (ML), a more severe clinical formof thedisease. Clinical aspect variability indicates that, besides the genetic host background, antigenic strain differences of this*Leishmania* speciesalso drive infection fates (8,9,10).

*L. braziliensis* promastigotes migrate to draining lymph nodes after the insect bite,leading to the following possible outcomes: 1) infection control and subject evolution toeitheroligo or asymptomatic (11, 12); 2) diseaseprogress to nodules (early-CL), which increase in size and ulcerate (late-CL)(13); 3) clinical lesion cure after anti-leishmanial therapy or spontaneous healing (7, 14, 15); 4) evolution to mucosal leishmaniasis in 3 to 5% of infected subjects (16). Pioneer studies applyingexperimental animal models (17, 18, 19) and, to a lesser amount, in humans (20, 21, 22, 23) have shown that the early infection phase is crucial in determining *Leishmania* infection fate. Infective burden (24), cytokine balance (25), infection site (26) and infection-site infiltrating leukocyte dynamics (27, 28) duringthe very early infection stageseemto dictate and determinethe evolution of the disease.

During the natural history of thisinfection, *L. (L.) major*-C57BL/6 modelparasites slowly multiply during the first 2 weeks, reaching a peak around the 4^th^ week post-infection (pi). This “silent phase” coincides with increases in IL-4 expression. IFN-γincreased from the 4^th^ week, coinciding with the appearance of nodular lesions (5^th^week) and decreasing parasiteloads (19). In BALB/c mice, rapid IL-4 production soon after *L. majorinfection* is necessary and sufficient to instruct Th2 cell development, resulting in disease progression(18, 25, 29, 30). However, little attention is given to the mechanisms underlying the first steps of the parasite-host relationship that influencedifferent *L. braziliensis* infection fates.

We have previously demonstratedthat hamstersinfected with 10^4^*L. braziliensis* promastigotes develop smallerlesions exhibiting less severe clinical aspects andtissue inflammatory infiltrate compared to animals infected by 10^5^or 10^6^inocula. Despite this, high IFN-γgene expression, anti-*Leishmania* IgG levels, and parasite load occurred independently of inocula size. However, noduleappearance onsetis indirectly correlatedto the number of inoculated parasites, indicatingthat parasite loads influence the clinical course of leishmaniasis (31).This prompted the investigation of early immunological profiles associated to favorable prognoses of infected animals exposed to lower parasite loads (10^4^) in comparison to those developing the severe form of disease, challenged with higher parasite inocula (10^5^ or 10^6^).Therefore, a study was designed in which an *L. braziliensis*-hamster outbred model challenged with three different parasite burden tracked at the main disease stages would expect to mirror the natural history of CL in humans.

## Results

### *Low* Leishmania (Viannia) braziliensis *infection is associated with a better clinical prognosis*

Lesion development kinetics(Figure 1B), confirmed a progressive course of evolution towards the chronic disease phase. An inverse relationship between parasitic inoculum concentration and lesion time onset was reproduced (Figure 1A) (31). At 50 dpi, the median lesion size in animals infected with 10^5^ parasites wassignificantly higher (1.00 mm [0.86 – 1.51mm]) when compared to the 10^4^group (0.24 mm [0.08 – 0.88 mm]), but lower than the 10^6^group (1.59 mm [1.39 – 1.93 mm]).

**Fig 1.**
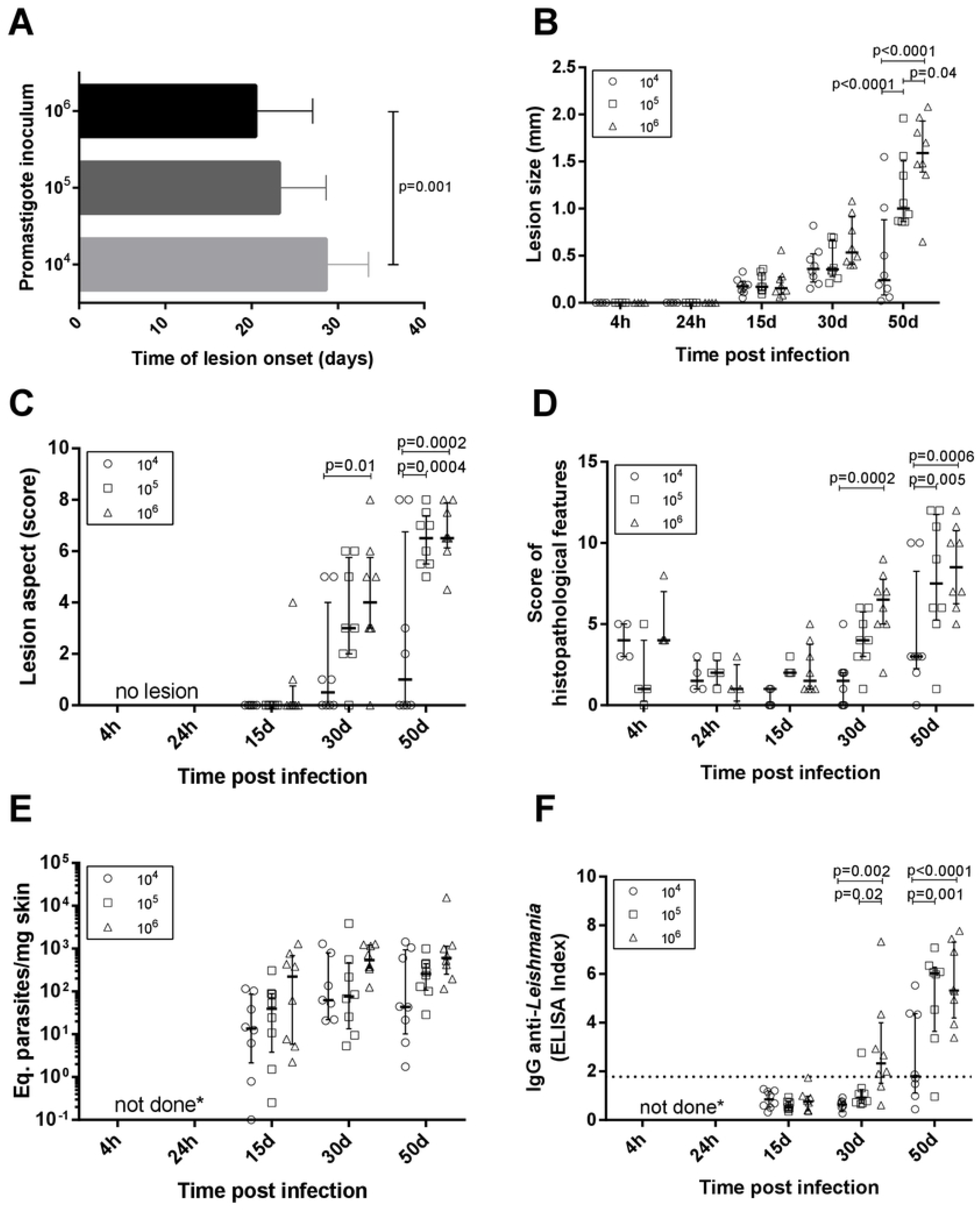
Clinical, parasitological and immunological aspects in golden hamstersinfectedwith *Leishmania (Viannia) braziliensis*. The animals were followed-up at during thefirst hours (4h, 24h), early (15 days and 30 days) and late (50 days) post-infection phases. A. Time of lesion onset; B: Lesion size was defined by the difference in millimeters (mm) between the thickness of the infected and the non-infected paw of the same animal; C. Lesions scores (s) were defined as follows: no lesion (s=0), edema (s=1), discrete papule (s=2), moderate papule (s=3), discrete nodule (s=4), moderate nodule (s=5), large nodule (s=6), very large nodule (s=7), and, if ulcerswere present, associated withearly ulcer (s=0.5), ulcer (s=1), ulcer with crust (s=1.5), deep ulcer (s=2);D. Regarding histopathological scores: no observation (score=0), slight (score=1), moderate (score=2) and intense (score=3); E. Parasite loadswere quantified in the lesions by qPCR and the results were expressed as Equivalent parasites per milligram of skin; E. Anti-IgG levels were quantified in plasma by an ELISA and the results were expressed as an Elisa Index. 10^4^ (o), 10^5^ (**□**), 10^6^(**□**)–number of promastigotes used for infection; Each symbol represents one animal; The lines in each scatter group represent the median and interquartile range; the dotted line indicatesthe cutoff value for IgG quantitation; *p* – Statistical significance.

A variable lesion increment pattern was observed, especially at 50 dpi. At this time point, most animals infected with 10^4^ parasites (six out of eight) presented less than 0.5 mm swelling. Two animals, however, exhibited lesions as large as those observed in the 10^5^ or 10^6^groups. On the other hand, one out of eight animals infected with 10^6^ parasites developed a smaller lesion (0.65 mm) than the median observed in this group (1.6 mm) (Figure 1B). Mostanimals in the10^4^grouppresentedless severe lesions than those infected with 10^5^ and 10^6^ parasites (Figure 1C). Curiously, despite differences in lesion size, these two groups presented similar clinical sign scores and the predominance of nodular and ulcerated lesions, considered a more destructive outcome. Similar patterns were observed concerning thehistopathological scores (Figure 1D).

### *Anti*-Leishmania *IgG is higher in animals infected with 10^5^ and 10^6^promastigote loads compared to 10^4^*

Results were expressed as the Elisa index (Figure 1F). No specific IgG production was observed in any of the three experimentalgroups at 15 dpi. At 30 dpi, most animals fromthe 10^4^ and 10^5^groupsmaintained IgG anti-*Leishmania* levels under the cutoff(except a single 10^5^ infected animal), whilemost infected by 10^6^ parasites already exhibited seroconverted IgG levels. At 50 dpi, IgG levels in the10^4^ group were significantly lower than those in the10^5^and 10^6^.Animals presenting IgG under the cutoff at 50 dpi were similar,exhibitinglesion size less than 1.0 mm (Figure 1B) and parasite load under 10^2^ (Figure 1E).

### *Arginase and iNOS expression donot determine parasite loads in hamsters infected with* Leishmania (Viannia) braziliensis

DNA parasite quantitation in lesions at 15 dpiwas similar in all groups(Figure 1E).At 50 dpi, it is noteworthy that animals from the 10^4^ group presented variable amounts of parasite loads. Instead, the 10^5^ and 10^6^ groups exhibitedsimilar patternsand lower variability in comparison to the 10^4^ group. Surprisingly, no parasites were observedin the histopathological analysis at 4 hpi. Immunohistochemistry assessments indicated diffuse staining throughout the dermis,but no integral parasites. At 24hpi, staining was more concentrated insebaceous glands and hair follicles and amastigote groups were identified (Figure 2). No inflammatory cells were observed.

**Fig 2.**
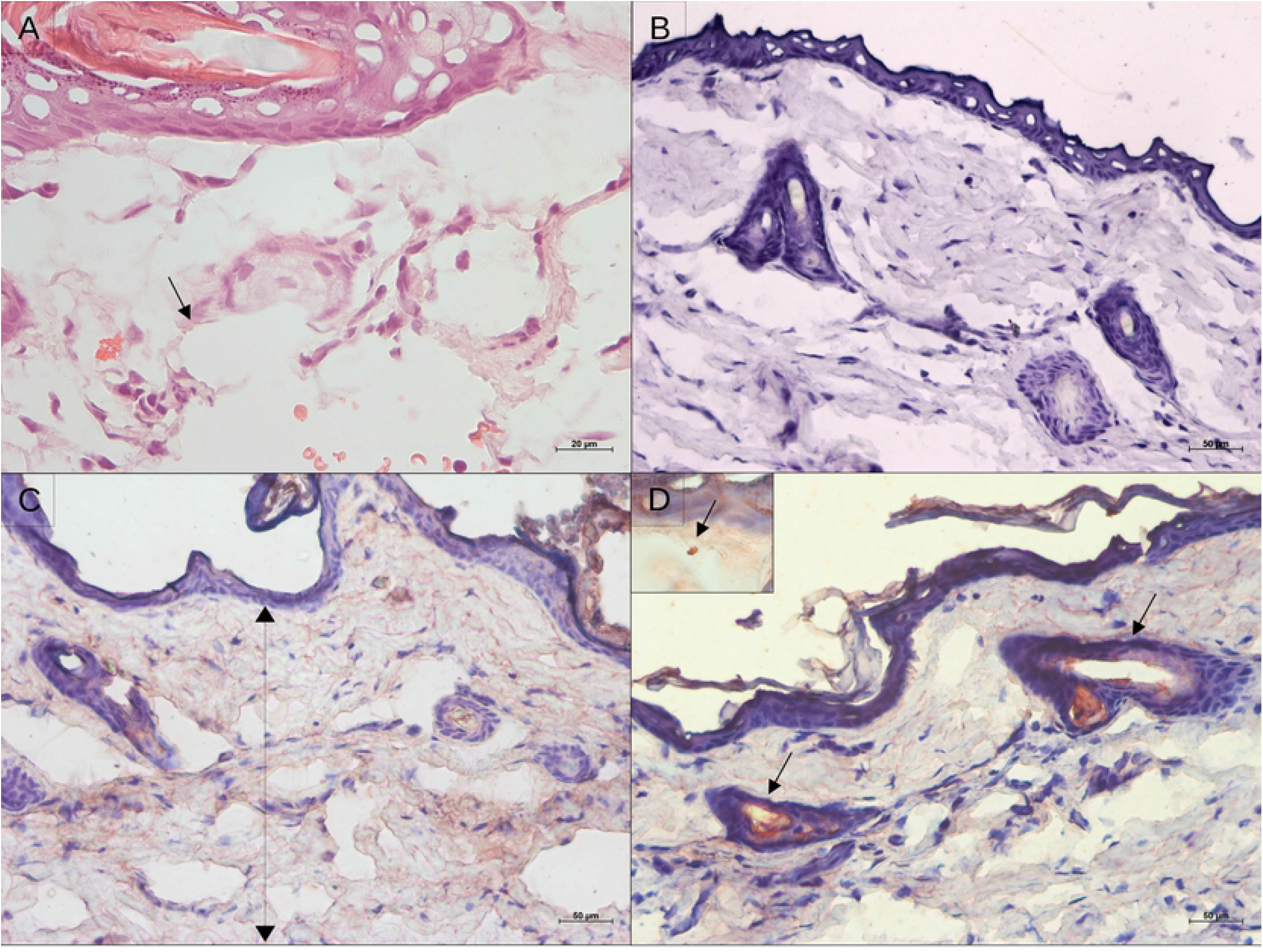
Histological aspect and leishmanial antigen detection in the skin of golden hamstersduring early after *Leishmania (Viannia) braziliensis* infection. A. Epidermal and dermal structure at 4 hours pos-infection, showing no tissue disturbance, only intumescent lymphatic vessels (hematoxylin and eosin; 20X magnification); B. Immunohistochemistry control; C. Immunostainning for *Leishmania* antigens showing antigensspread throughout the tissue at 4h, and D.Detection of leishmanial antigens at 24hpi showing concentrated staining of the nearest pilous follicle (50X magnification). The insert displays an amastigote.

Arginase, an enzyme found in all *Leishmania* strains and produced by macrophages, stimulatesparasite replicationand inhibitsnitric oxide function (32). The basal *arg* expression was high in hamster skin. At 4 hpi,*arg* expression was upregulated compared to normal skinin the 10^5^ and 10^6^groups, but not in the 10^4^ group. After 24 hpi,an *arg* increase was also observedin the 10^4^ group. All three inocula groups exhibitedaround a 5-foldincrease in*arg* expressionat 15 and 30dpi, and around a 10-foldincreaseat 50 dpi (Figure 3A and 4A). This environment is accompanied by elevated parasite loads (Figure E).

**Fig 3.**
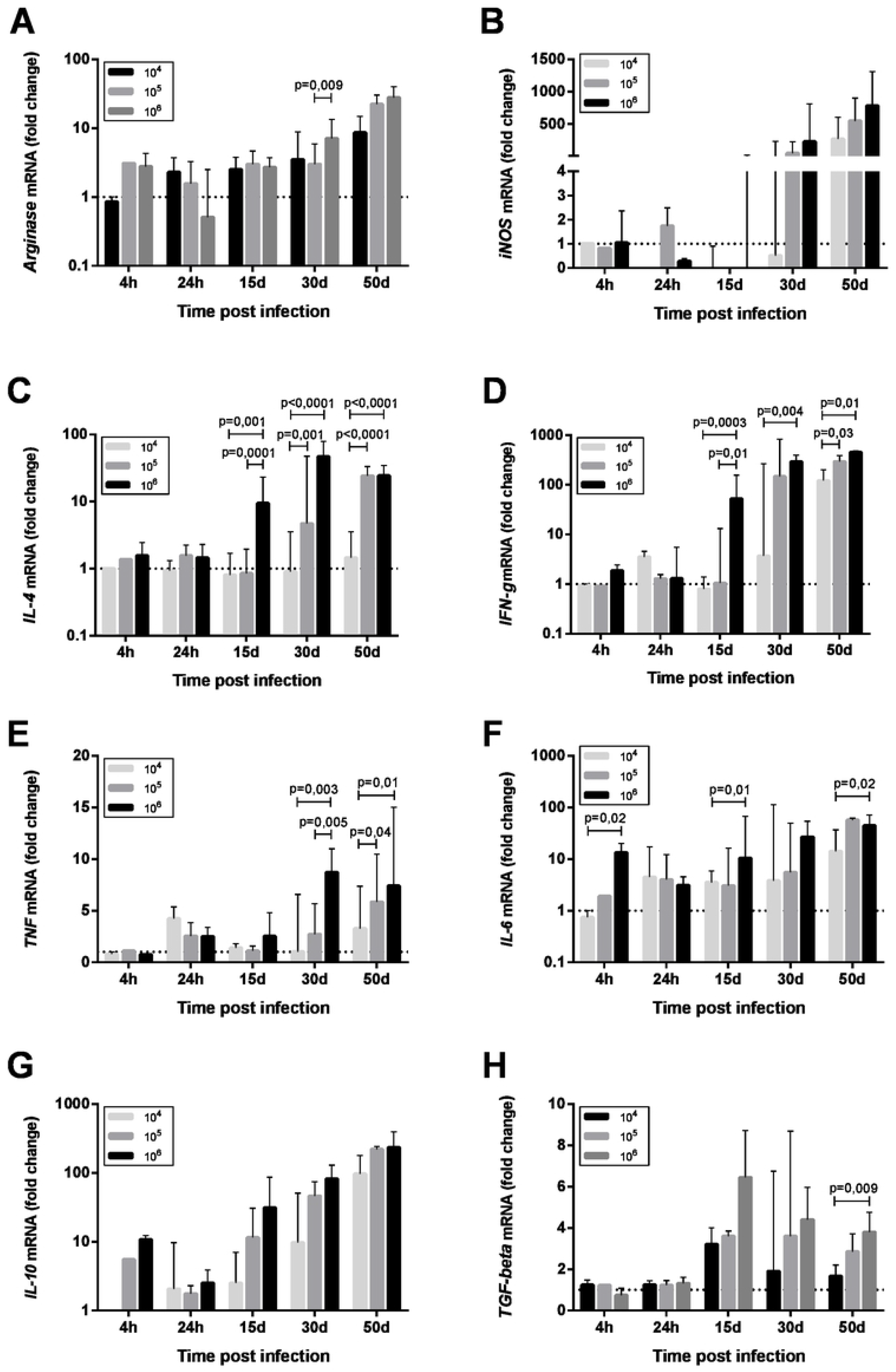
Cytokine mRNA expression in the skin of golden hamsters at different time points after infection with different *Leishmania* (*Viannia*) *braziliensis*. The animals were infected with three parasite concentrations (10^4^, 10^5^, 10^6^ promastigotes). Skin mRNA expression was determined by RT-qPCR at different time post infection, namely thefirst hours (4h, 24h), early (15 days and 30 days) and late (50 days) post-infection phases. The dotted line represents the basal expression of non-infected skin. The results are expressed as relative fold changes between the experimental samples and the skin of an uninfected animal, to which a value 1 was arbitrarily assigned. The bar represents the median and the line, the interquartile ranges. 10^4^, 10^5^, 10^6^ –number of promastigotes used for infection.

**Fig 4.**
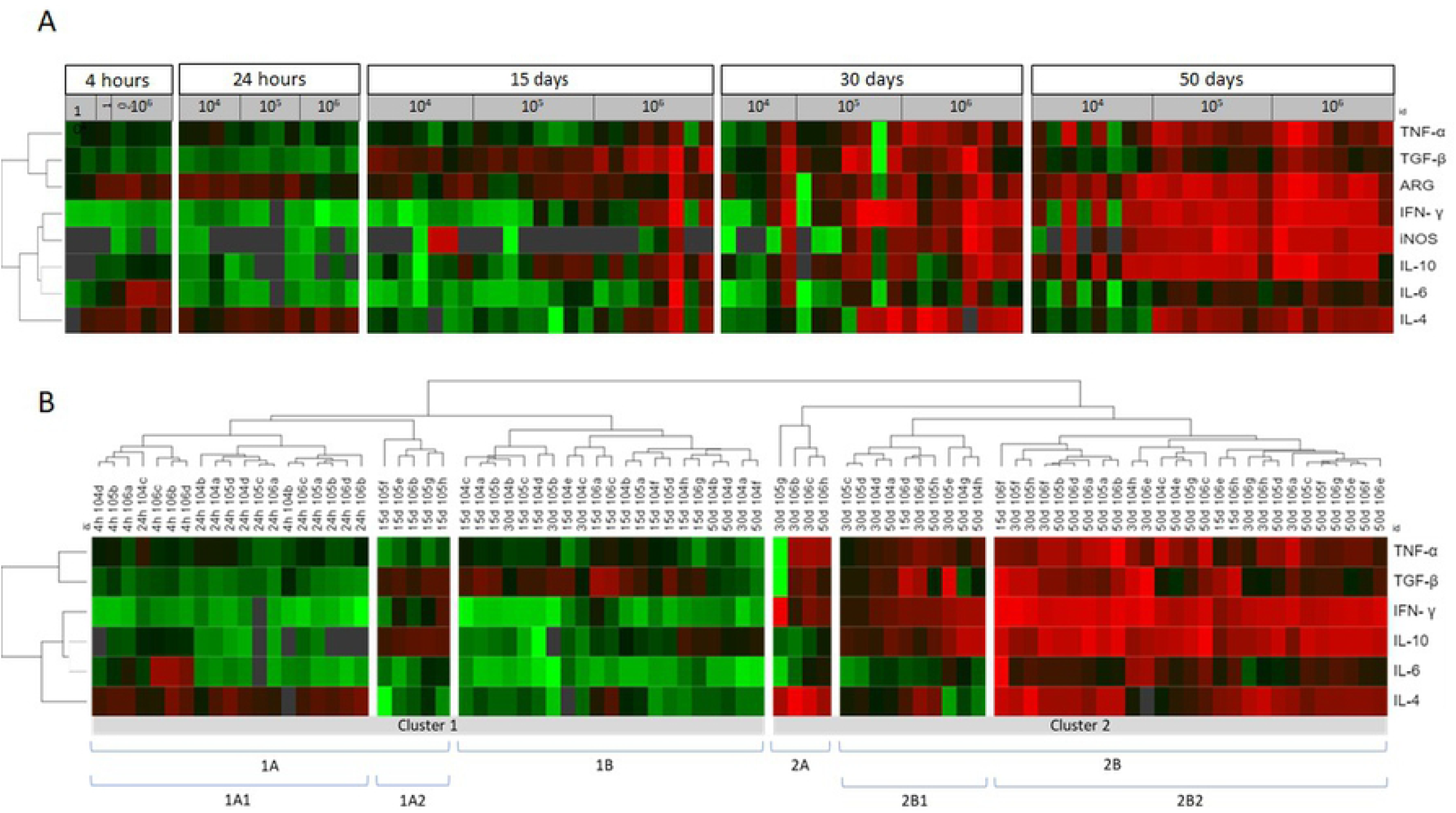
Cytokine, iNOS and arginase in the skin lesions of hamsters infected with *Leishmania (Viannia) braziliensis* according to a comparative gene expression heatmap analysis. (**A**) The animalswere grouped by post infection time (4 and 24 hours; and 15, 30 and 50 days) anddifferentinocula (10^4^, 10^5^, and 10^6^ promastigotes). Heatmap analyses were performed for gene expression (*tnf, tgfβ, ifnγ, il10, il6, il4, arg, inos*)ΔΔCt using the online Heat mapper® software. (**B**) The animalswere clustered by gene expression signature. The hierarchyaverage linkage clustering method withEuclidean distance measurement was used (Wishart Research Group at the University of Alberta). Each row represents one molecule and each column represents one animal. Higher gene expression is displayed in red,and lower gene gene expression, in green. Gray indicates mean gene expression below the lower limit of detection.

Macrophage microbicidal function was evaluated byiNOS gene expression (Figure 3B and 4A). Basal iNOS expression in hamster skin is low and was down-modulated or even absent (gray color) duringthe first hours (4- and 24-hpi), as well as at 15 dpi, independent of the inocula. At 30 dpi, a similar trendwas observed for animals from the 10^4^ group and in half of the animals infected with 10^5^, whilethe 10^6^ group exhibiteda high expression of this enzyme. At 50 dpi, high iNOS expressionsweredetected in half of the eight animals from the 10^4^ group and in all the animals from 10^5^ and 10^6^ groups. However, thisiNOs expression increasewas not associated with parasite load control (33).

### Promastigote infective loads dictate the parasite-induced cytokine pattern

Changes in thecytokine gene expression of infected skin compared to uninfected skin were assessed. At the moment of infection,the promastigotesencountered an environmentof high TGF-βgene expressionand intermediary basal TNF, IL-6, and IL-4 geneexpressions. Concerningthe 10^4^ group,promastigotes did not trigger an increase inIL-4 expression at 4 hpi (Figure 3). Basal IL-4 levels were maintained up to 30 dpi, and a discrete upregulation was observed only at 50 dpi (Figure 3C). On the other hand, an IFN-γ and TNF wave was observed at 24 hpi(Figure 3D and 3E). IL-6 and IL-10 were also upregulated at 24 hpiin the 10^4^ group, and were maintained or even increased up to 50 dpi (Figure 3F and 3G). TGF-βupregulation was observed from 15 dpi to 50 dpi (Figure 3H).

In contrast, a slight IL-4 trigger was observedsoon after the infectionin the 10^6^ group (Figure 3C), while IL-6and IL-10 increasedaround twoto 10-fold (Figure 3F and 3G). In addition to TGF-β, TNF and IFN-γ, these cytokine genesexhibited highexpression maintenance during the early phase time points (15 and 30 dpi) and duringthe chronic phase when infected with 10^6^ (Figure 3H, 3E and 3D). The 10^5^ group exhibited a tendency to follow the 10^6^ group pattern, althoughthe alterations took longer to occur. A hallmark of the three experimental groups was the over-expression of all cytokine genes duringthe chronic phase. However, IL-4 expression intensity was much lower in the 10^4^groupin comparison to the 10^5^ and 10^6^ group (Figure 3C). These differencesprompted us to investigate possible cytokine genes profiles associated to the disease outcome.

### *IFN-*γ *gene expression and IgG anti-* Leishmania*influence lesion progression, while IL-6isrelated to parasite loads*

The factors underlying the clinical outcome of disease were also analyzed (Table), taking into account dependent (anti-*Leishmania* IgG, cytokines, arginase and iNOS gene expression) and independent (lesion size and skin parasite loads) variables. The applied multiple linear regression model was significant for both independent variables (p<0.05) and explained 66.7% (R^2^=0.667) of the total variance of lesion size and 62.6% (R2=0.626) of the parasite load. IFN-γ gene expression (coefficient β= 0.576; p=0.017)and IgG anti-*Leishmania*(coefficient β =0.375; p=0.007) werepositively correlated with lesion size. IL-6 was the only cytokine influencing parasite loads(coefficient β= 0.447; p=0.028).

### Cytokine gene signatures maybe related tolower parasite loadbut not to disease progression

We used heatmaps in an attemptto group animals presenting similar skin cytokine gene expression patterns (Figure 4A). The animals were clustered into 1A1, 1A2, 1B, 2A, 2B1 and 2B2, according to cytokine gene patterns (Figure 4B). Cluster 1A1 grouped the animals at 4- and 24hpi, without any segregation related to the differentinocula.The 1A2 and 1B clusters grouped most animals at 15 dpi (19 out 23), four animals at 30 dpi (threefrom the 10^4^group and one from the 10^5^group), and three50 dpi animals from the 10^4^ group. Even considering this heterogeneity, animals inboth clusters presented only discrete footpad swelling (Figure 5A and 5B) anddiscrete inflammatory infiltrates (Figure 5C). All other animals except for one presented aparasite load equal or lower than 10^2^Eq/skin/mg, including those at 30 or 50 dpi (Figure 5D).

**Fig 5.**
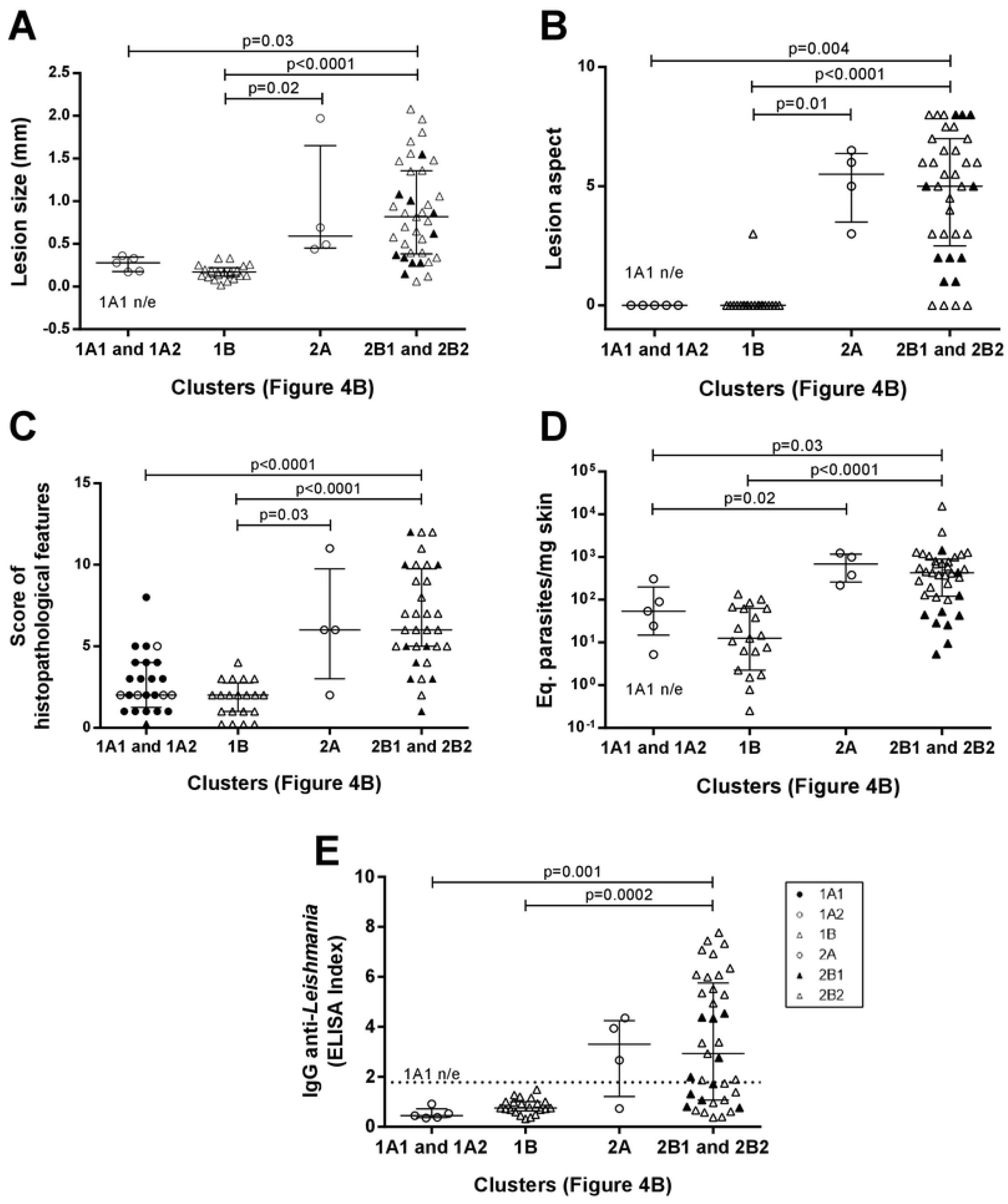
Clinical, parasitological and immunological aspects of *Leishmania (Viannia) braziliensis* hamster infection clustered according to skin cytokinegene expression signatures. The hierarchy average linkage clustering method with Euclidean distance measurement was used (Wishart Research Group at the University of Alberta) based on heat map analysis, as shown in Figure 4B (clusters: 1A1, 1A2, 1B, 2A, 2B1, and 2B2);Biomarker gene expression (*tnf, tgfβ, ifn*γ, *il10, il6, il4, arg, inos*); Each symbol represents one animal; The lines in each scatter group represent the median and interquartile range; the dotted line indicates the cutoff value for IgG quantitation; ▴ represent the 2B1 subcluster and • represent the 1A1 subcluster. *p*-statistical significance;

The 1A1 cluster was characterized by the up-modulation of theIL-4 gene (Figure 4B). In contrast, animals in the 1A2cluster exhibitedupregulatedTGF-β and IL-10 expressions andslightly downmodulated IL-4 expressions. Concerning1Bcluster,most animals presentedTGF-β upregulation. Four animals in this cluster (one at 30 dpi and three at 50 dpi, all from 10^4^ group) exhibiting low TGF-β presented high IL-10 expressions. Therefore, at least two cytokine gene profilesseem to definetheseclusterings. The 1A2 cytokine signature maybe associated with higher parasite loads (albeit,non-significantly) when compared to 1B (Figure 5D). On the other hand, greater parasite load variability was observed in the 1B group.

Cluster 2, especially 2B2, grouped mostanimals inthe late phase of the infection (50 dpi) or infected with higher inocula (Figure 4B). As expected, these animals presented more severe lesions, higher IgG anti-*Leishmania* and higher parasite loadsthanthose in cluster 1 (Figure 5). Aconsiderable variability ofall parameters, however, was observed. Of utmost importance, parasite load was the main distinction between subclusters 2B1 and 2B2. The 2B1 animals (black triangle)displayed significantly lower parasite loads than those in the2B2cluster (p=0.0005) (Figure 5D). The differencebetween these subclusters was the downregulation ofall cytokine expressions, more pronounced for IL-6in the 2B1 cluster when compared to the 2B2 group(Figure 4B). It is important to note that these cytokine signatures do not define animmune response pattern associated with disease control during thelatter infection phase (Figures 5A, 5B and 5C).

### An intensenetwork interaction is sustained in animals able to control the disease

To globally evaluate the clinical immunopathological status of the disease, a network analysis of the correlation matrix including all studied biological parameters (Figure 6) was carried out. The 10^4^ group presented the lowest level of interactions at 15 dpi in comparison to the other groups. The number of interactions increased overtime in the 10^4^ group, contrasting to the scarce number of interactionsobserved in the 10^5^ and 10^6^ groups. At 50 dpi, the 10^4^ group exhibitednumerous strong positive correlations involving IFN-γ and regulatory cytokines (IL-10 e TGF-β), as well as TNF/IL-6 with anti-inflammatory cytokines (IL-4). This pattern was associated with animals presenting partial parasite load (Figure 1E) or lesion size (Figure 1B) control.

**Fig 6.**
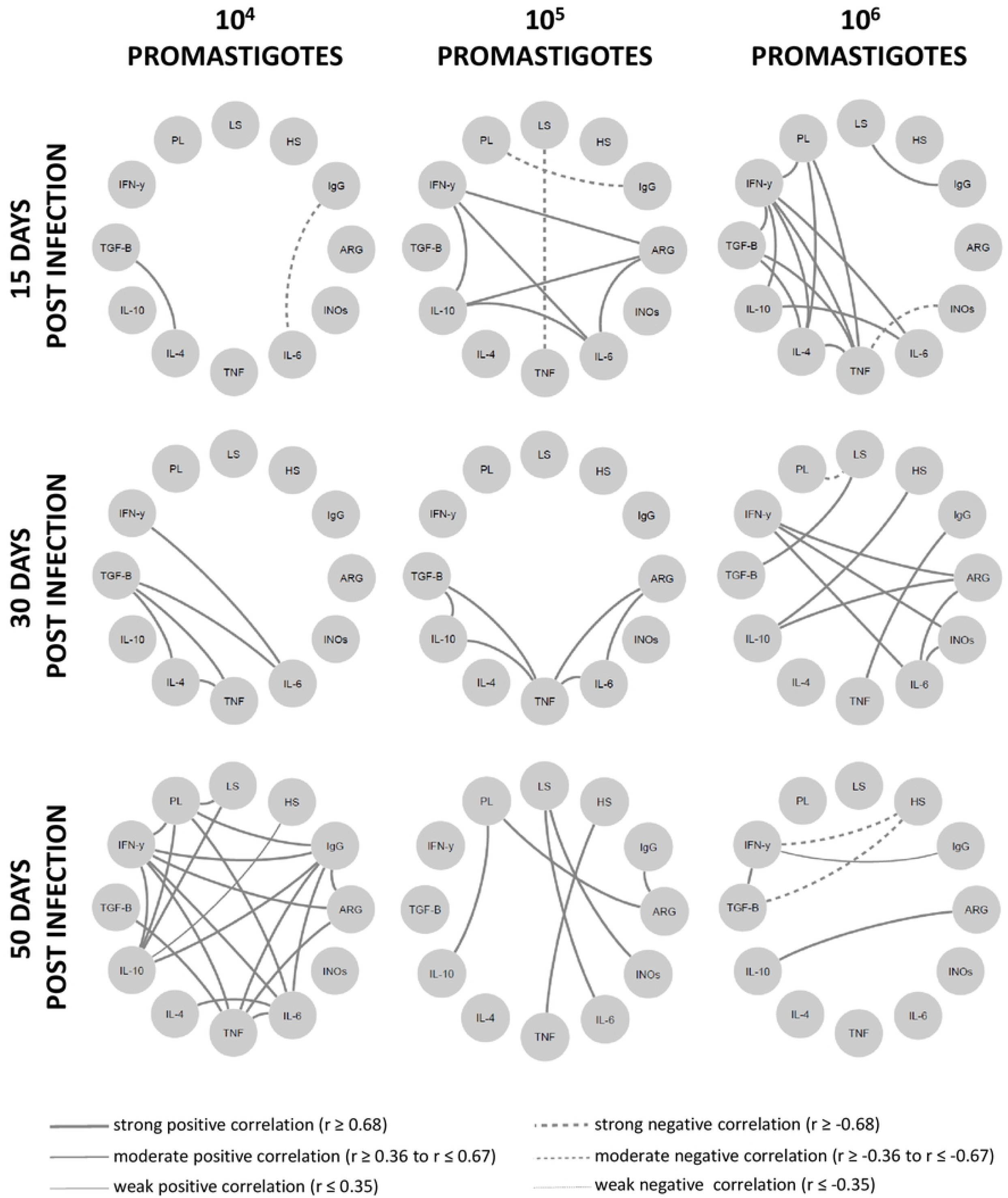
Network interaction involving clinical, parasitological and immunological biomarkers in hamsters infected with different *Leishmania Vianna braziliensis* inocula. Hamsterswere evaluated during the first hours (4h, 24h), early (15 days and 30 days) and phases (50 days) post infection phases with 10^4^, 10^5^ or 10^6^ promastigotes. Significant Spearman correlationswere considered when *p* ≤0.05, represented by lines connecting the circle nodes. The correlation index strength was represented by the connecting line thicknesses. PL - parasite load; LS – lesion size; HS – histopathology score; ARG-arginase; INOs – Oxid nitric sintase; Cytokine genes – TNF, TGF-*β*, IFN-γ, IL-10, IL-6, IL-4.

To certify whether the disease control was associated with an intense network interaction, we distributed 50 dpi animals into two groups, namely the disease control group (lesions smaller than 1mm and parasite load <10^2^); and the disease progression group (lesions largerthan 1mm and parasite load > 10^2^). The signature observed at 50 dpi inthe 10^4^ group was reproduced by the disease control group,exhibiting even more intense strong and positive interactions (Figure 7). Onthe other hand, the disease progression group presented a network displayinga lower number of moderate and positive or moderate and negative interactions, sustaining the hypothesis that immunopathogenesis is associated toadownmodulated immune response.

**Fig 7.**
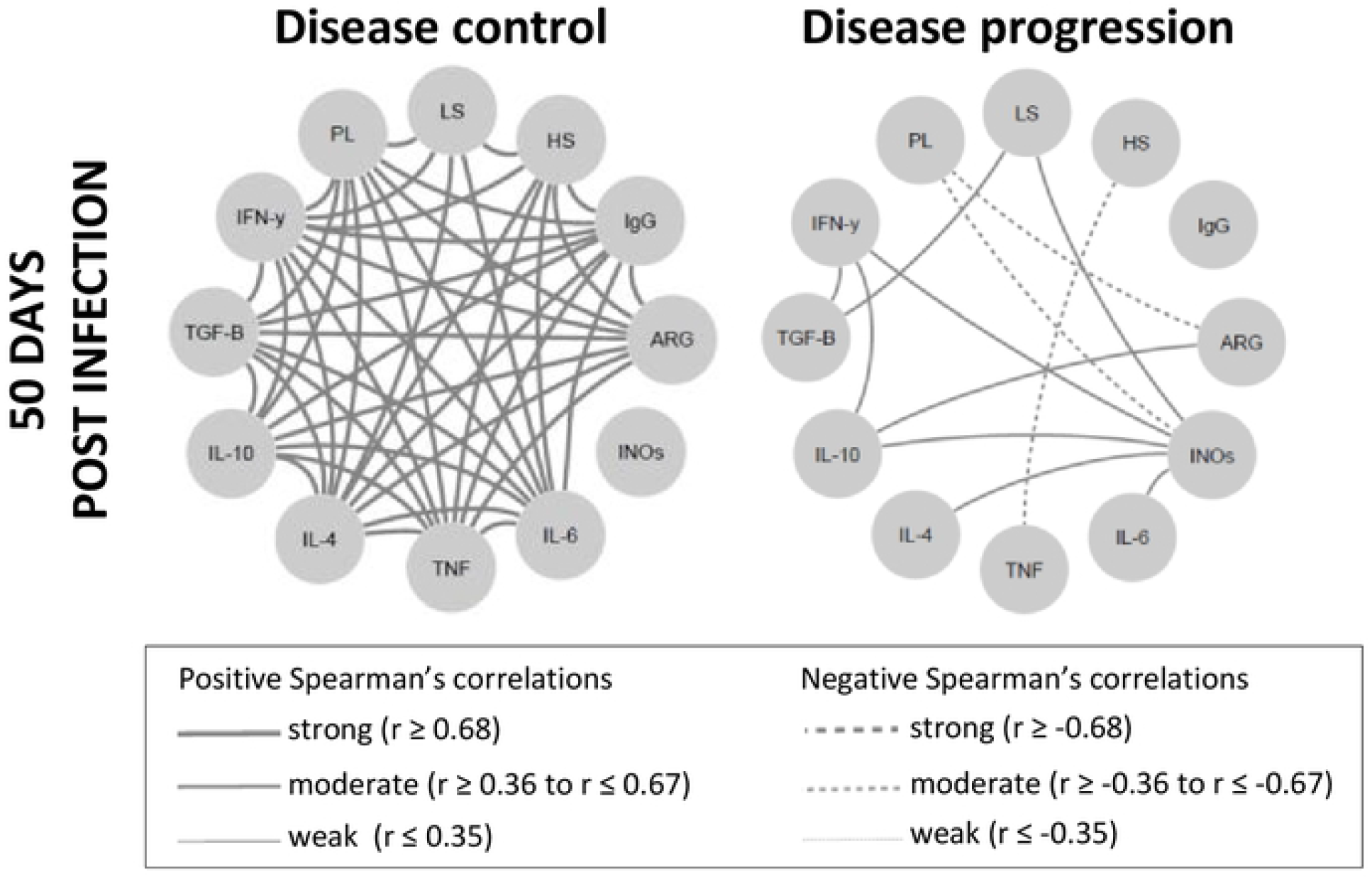
Skin lesion clinical outcome is correlated with degree of network interaction involving clinical, parasitological and immunological biomarkers in *Leishmania Vianna braziliensis*-infected hamsters. Network interactions evaluated between animals were subdivided into two clinical fates, according to lesion size and parasite load, asdisease control (lesions smaller than 1mm and parasitic load <10^2^)and disease progression (lesions larger than 1mm and parasitic load > 10^2^). Significant Spearman correlations were considered when *p* ≤0.05and were represented by lines connecting the circle nodes. The strength of the correlation index was represented by the connecting line thicknesses. PL – parasite load; LS – lesion size; HS – histopathology score; ARG-arginase; INOs –oxid nitric sintase; Cytokine genes – TNF, TGF-*β*, IFN-γ, IL-10, IL-6, IL-4.

## Discussion

Duringnatural *L. major* infection, it is suggested that a sand fly transfers<600 promastigotes, although higher inocula up to 100,000 have been reported (34), as observed for other *Leishmania* species (35). The infectious load can influence the severity of leishmaniasis, as higher inocula accelerate the pathogenic process (24, 31, 34), whereas lower parasite loadsseem to be associated with infection self-control, a common outcome in endemic areas (36).

At the infection, the promastigotes encountered a favorable skin environment to replicate, with high arginase and TGF-β,as well asan intermediary IL-4 expression and low iNOS. However, most promastigotes were probably destroyed *in situ*, as suggested by the detection of *Leishmania* antigens scattered throughoutthe dermis, suggesting innate immunity action without disturbing tissue architecture. The interaction betweenpromastigotes and theskin immune system during thevery short time before their entry into host cells results in local cell stimulation. We observed that IL-10 and IL-6 genes were induced in the skin of animals infected with 10^5^ and 10^6^ inocula, while no changes were observed for TGF-β. Curiously, IL-6 positively influenced the parasite loadvariable. WhetherIL-6 acts directly on parasite growth or via anti-inflammatory responses as a consequence of classical membrane receptor signaling (37) must still be clarified. PBMC fromhealthy humans stimulated by promastigotes also increased IL-10 production, but not IL-6, while TGF-β was down-modulated (38). It is possible that the number of parasites challenging the immune system drives these phenomena, as no cytokine induction was observed in the 10^4^ group at 4 hpi. Very few amastigotes inside cells were observedat 24 hpi, as also reported elsewhere(28). On the other hand, it is important to notethat only the 10^4^ group evaluated at 24 hpi presented TNF and IFN-γ geneupregulation, potentiallyassociatedto the recently transformed amastigotes. This wave could have exerted apositive influence in the next infection steps.These findings indicate that the infective burden influences parasite-driven cytokine activation, in which IL-4, IL-10 and IL-6 activation seems to playan infection progression role.

IL-4 gene induction was also differentamong the threeinfected animalgroups. A slight IL-4 gene upregulation was observed at 4 and 24 hpi in the 10^5^ and 10^6^ groups, but not in the 10^4^ group. Alongsideno TGF-β, IL-10 or IL-6 gene upregulation in the 10^4^ group, this points to a less permissive parasite proliferation environment. On the other hand, an intermediary basal expression of the IL-4 gene in hamster skin was confirmed (31). This cytokine hasalready been shown to be produced by activated skin mast cells (39, 40) or keratinocytes (41). Early IL-4 expression is noteworthy as an important resident dendritic cell activation and differentiation mediator (42, 43). Interestingly, this phenomenon was also demonstrated as a requirementfor Th1 response activation (42, 44).

Although no clinical lesion signswere observed until 15 dpi, a considerable number of parasites were already detected in the lesion sites. This fits with the “silent phase” described in the CD57/BL6-*L. major* model, in which establishment of apeak load of parasites in the dermis occurred in the absence of lesion formation (19)or any overt histopathological*in situ* alterations. In humans, parasitized lymphoadenopathy also precedes CL lesions (45). Herein, the parasite burden also increased through the assessed time points,indicatingongoing replication, despite high iNOS expressions, indicating no microbicidal effect (33).

After the silent phase, a TGF-β upregulation at 15 dpi was noted, when the effects of a *Leishmania*-induced T-cell response take place. At this time, the lesions became apparent and an increase in macrophage infiltrates was observed. In the *L. (V.) panamensis-hamster* model, these cells have also been reported as appearingafter 10 dpi (28) to 15 dpi (46). In Balb/c macrophages, TGF-β augmentsthe replication of *L. braziliensis* which, in turn, also producesits biologically active product (47). These authors also observed an *in vivo* exacerbation of leishmaniasis lesions, and associatedIL-10, but not IL-4, induction to immune response downregulation. Corroborating this, *L. braziliensis* amastigote stimulation increased TGF-β induction 10-foldin human healthy donors, as well as IL-10, IL-6 and IL-1β mRNA, although less intensely. These authors also detected cytokine releasesinthese stimulated PMBC (38). It has been previously reported that six*L. braziliensis* strains induce different inflammatory and regulatory cytokines genes at 30 and 60 dpi, withlower gene induction observed for IL-6 (48). Curiously, IL-6 was the only dependent variable positively correlated with parasite load in the present study. As we utilized the same batch of a single*L. braziliensis* strain, the influence of host genetic background heterogeneity on the infection outcome was also clearly observed.

The lowest inoculum was most frequently associated with less severeskin lesions.At 50 dpi, the lesions were small or even absentin most animals of the 10^4^ group, with similar parasite loadsin the 10^5^ and 10^6^ groups. Both the 10^5^and10^6^ groups presented similar lesions aspects, histological damage, parasite load *andanti-Leishmania* IgG antibodies. This, however, didnot occur in all animals, resulting in a very heterogeneous clinical and parasitological profile. As previously reported(31), the 10^5^ group exhibited a similar trend as the 10^6^group after the early phase. This is noteworthy,considering the highdifference of theinitial inocula (900.000 promastigotes), and similar behavior between the 10^5^ and 10^6^ groups was unexpected. Curiously, CL lesion parasite loadsin humansdonot seem to be a good predictor of disease progression (49). The mechanism driving the clinical evolution of the 10^5^ group toward to 10^6^one seems to occur at the final early phase of the infection. The delay in disease development observed in the 10^5^ group suggest that anti-leishmanial immune mechanisms were initially triggered but were not sufficient to overwhelm the parasite burden. Other lesion analysis tools, such as changes in thespectral signature recently described for the *L. braziliensis* hamster model, will extend our understandingof skinformation and development (50).An up-regulation of almost all investigated cytokines was observed duringthe chronic phase. However, due toslight differences in their expressions, we considered that a systemic analysis could shedlight on this phenomenon.

The heatmap clusterization according to cytokine expression intensity clearly separated animals according to disease evolution time. All animals evaluated at 4- and 24 hpi (CL 1A1) and mostassessed at 15 and 30 dpi (CL 1A2 and 1B) were grouped. On the other hand, the CL2 group included, mainly, the animals at the end of the early and late infection phases. This clusterpresented a higher cytokine expression in comparison to CL1. The exception was for IL-4,whichpresented much lower relative quantitation in the 10^4^ group compared to the 10^5^ and 10^6^ groups.Meanwhile, the multivariate analysis demonstratedthat the IFN-γ gene, which was highly expressed, was the only independent variable that directly influenceslesion size, strengthening the role of imbalanced proinflammatory responseson CL pathogenesis (51, 52, 53). Although no cytokine influenced parasite loads, CL 2B1 presented lower parasite loads than CL 2B2. Interestingly, IL-6 gene expression was also differentially expressed in these clusters, comparativelyless expressed in CL 2B1 in comparison to CL2B2. This reinforcesthe potential role of IL-6 andIL-4 in parasite replication in the applied hamstermodel.On the other hand, IL-4 induction can also be associatedto a Th2 response that contributes to CL progression (30).

Cytokine inter-regulation has beendemonstrated in human leishmaniasis caused by *L. braziliensis* (51). These authorsobserved that IL-10 and TGF-β downregulated TNF and IL-17 production, whereas IFN-γ and TNF positively induced each other. On the other hand, neutralization of IFN-γ, but not TNF, induced IL-10 release. Thisis in agreement with the hypothesis that acytokine balance rather than their amountsaffect the clinical outcome. The network analysis conducted herein concerning theclinical and expression of genes involved in the pathogenesis indicated thathamsterswith the ability to control the *L. braziliensis* disease at 50 dpi adapted their immune response to a lower general cytokine expression coupled to a cytokine expression balance. These conditions probably maintained the regulation of cytokine effector functions through the production of relative amounts of soluble antagonistic factors. This harmonic signature involving the cytokines observed in animals that were able to controlCL maybe translated by the intense network of numerous interactions based on strong and positive Spearman correlations.The absence of negative correlations also reinforces the idea of a well-modulated immune response (54, 55, 56). On the other hand, animals exhibitingdisease progression displayeda reduced number cytokine expression relationships, establishing a low immunomodulation environment.Moreover, an inverse correlation betweenIFN-γ gene expressionanddisease progression alongside an unwavering IL-4 expressionthroughout the evaluated timeframe, were also considered asdefining animmunological hamster signature associated with CL control.

Takentogether, our findings indicatethat a lower infective burden results ina cytokine profile that favorsless severe cutaneous lesionsin the *L. braziliensis*-hamster model,althoughindividual susceptibility can subvert these protective mechanisms. IL-4, IL-6, IL-10, and TGF-β were associated with infection progression, while IFN-γ was corelated to tissue damage. However, more than the effect of a specific cytokine, the fine interrelation among cytokineswill dictate the clinical fate of *L. braziliensis* infection. This feature should be strongly considered invaccine designs.

## Material and methods

### Animals and ethics statements

Adult female outbred golden hamsters (*M. auratus*) (6-10 weeks old), weighing 80-90 g, obtained from the animal facilities belonging tothe Fundação Oswaldo Cruz (FIOCRUZ), were used. Ninety-six (96) infected animals(four/group) were included: a) one experiment for 4- and 24 hpi; b) two experiments for 15-, 30- and 50 dpi. Sixteen uninfected animals were used as control. This study was performed in two independent experiments and approved by the Ethics Committee on Animal Use (CEUA) of FIOCRUZ, with protocol number IOC 032/15.

### Parasites, infection and experimental protocol

*L. (V.) braziliensis* (MCAN/BR/98/R619) in the stationary growth phase from the third *in vitro* passage were ressuspended in a total volume of 20 μL of phosphate-buffered saline (PBS). Inocula containing 1×10^4^, 1×10^5^ or 1×10^6^ parasites were used for intradermal inoculation into the dorsal hind paw of hamsters, as described (31). Animals were euthanized 4- and 24-hours post-infection (hpi) and 15-, 30- and 50-days post-infection (dpi) through administration of the preanesthetic medications Ketamine and Xylazine (200- and −10mg/kg, respectively) injected intraperitoneally and, after sedation, sodiumthiopental at a dose of 150 mg/kg, also intraperitoneally. The death of the animals was confirmed by cardiac arrest.

### Clinical evaluation of *Leishmania (Viannia) braziliensis* infection

The skin lesions were monitored weekly from day 7 up to 50 dpi, measuring the paw dorsum-ventral thickness with a digital thickness gauge (Mitutoyo America Corporation, São Paulo, Brazil). The lesion size was determined by the difference in millimeters between the thickness of the infected and the non-infected paw of the same animal. Clinical aspects of the lesions were qualified with a score system (Figure 1C).

### Quantification of *anti-Leishmania* antibodies

The levels of IgG *anti-Leishmania* were determined by an ELISA assay in house (Enzyme-linked immunosorbent assay) as previously described with some adaptations (57). Briefly, *L. braziliensis* (MHOM/BR/1975/M2903) soluble antigen (2 μg/well) were adsorbed in microtiter plate (Nunc-immuno Plate, Roskilde, Denmark). Then, hamsters’ plasma samples were diluted 1:50 and added in triplicate. Horseradish peroxidase labelled goat anti-hamster IgG was used as detector system (Santa Cruz Biotechnology, Santa Cruz, CA, USA). The results were expressed as ELISA index (EI), obtained by mean of sample absorbance divided by mean of negative controls (n=5) absorbance. The cut off was determined by ROC method (Receiver Operator Characteristic Curve).

### Histopathological analyses

Fragments from the skin of the infected paw were fixed in 4% paraformaldehyde solution and processed for paraffin embedding. Sections of 5 μM thickness were stained with haematoxylin-eosin and then observed by light microscopy (Nikon Eclipse E600, Microscope, Tokyo, Japan). Results from the skin histopathological analysis were expressed by a score criteria (31), based on a semi-quantitative analysis that evaluated the intensity of each histopathological features occurred: presence of inflammatory infiltrated, bleeding, congestion of vessels, granulomas, vacuolated macrophages, *Leishmania* amastigotes, Schaumann’s bodies and/or necrosis (Figure 1D).

### Parasites detection by immunohistochemistry

Another skin fragment was embedded in OCT compound (ornithine carbamyl transferase compound; Tissue Tek, Illinois, USA). Frozen tissue sections (4 μM) were cut at −27°C and mounted on poly-L-lysine (Sigma Chemical Co., Saint Louis, USA) coated slides. The slides were dried, fixed in cold acetone (Merck, Darmstadt, DE) for 10 minutes and permeabilize with 0.4% Triton X-100 (Sigma, USA) in Tris-HCl. Then, samples were incubated with 0,6% hydrogen peroxide to block endogenous peroxidases and were then incubated with 2% bovine serum albumin (Sigma, USA) to inhibit non-specific binding. Heterologous serum (primary antibody) collected from mice infected with *L. (L.) infantum* (IFLA/BR/1967/PH8) was incubated overnight at 4°C in humidified chamber diluted 1:100. The Polymer Detection System for Human Tissue (Abcam, Cambridge, UK) was used for immunohistochemical staining following the manufacturer’s instructions. The reaction was revealed using aminoethyl carbazole as chromogen. Nuclei were counterstained with Mayer’s hematoxylin (Merck), and the slides were mounted in aqueous mounting medium. Normal mouse or goat serum was used as negative control.

### Cytokine, arginase and iNOS gene expression by reverse transcription quantitative PCR (RT-qPCR)

Cytokines and enzymes gene expression were quantitated from skin lesion fragment using a combination of TRIzol® (Invitrogen, California, USA) and RNeasy® minikit (Qiagen, Austin, Texas, USA) for RNA extraction (water phase). The evaluation of IFN-γ, TNF, IL-6, IL-10, TGF-β, IL-4, iNOS, arginase and housekeeping GAPDH and γActin mRNA were based on the protocol previously described (58). Relative quantitation of gene expression was calculated using the comparative Ct method (ΔΔCt), as previously described (59), with threshold set at 0.02. Gene expression was represented as fold change (2-ΔΔCt), in relation to skin samples from uninfected hamsters, used as calibrators. The Expression Suite Software (Thermo Fisher Scientific, Massachusetts, USA) was used for analysis.

### Parasite load detection by quantitative PCR (qPCR)

Parasite load was quantified from the same skin lesion fragment used to determine mRNA expression, using the TRIzol® protocol for DNA extraction (organic phase). The measurement of parasite load by qPCR was performed by absolute quantitation, based on a standard curve produced from DNA samples extracted from fragments of hamster skin, artificially infected with promastigote forms of *L. braziliensis*. For parasite quantitation, primers targeting the conserved regions of kinetoplastid DNA minicircles (kDNA) were used (60). The parasite load was calculated by the (*L. braziliensis* equivalents/hamster skin mass equivalents) ratio and expressed as “parasite load” (parasites eq/mg skin).

### Statistical analyses

Comparison between experimental groups was performed using one-way analysis of variance (ANOVA). Statistical tests from the ΔCt values were student t-test or Mann-Whitney rank sum test, and Analysis of Variance. Spearman’s correlation matrix was performed using ΔCt values for gene expression. were performed with GraphPad Prism software version 6.0 for Windows (GraphPad Software, San Diego, CA, USA) or SigmaPlot v12.0 software (Systat Software, Inc). A multivariate statistical analysis was performed through multiple linear regression (SPSS software, version 9.0) to determine the influence of intervening variables. Heatmap matrix analyses were performed for gene expression ΔΔCt using online software Heat mapper® (Wishart Research Group at the University of Alberta). The hierarchical clustering method used for analysis was the average linkage, and the distance measurement method applied was Euclidean. The interaction network was done with correlations that presented significance level ≤0.05 using Cytoscape 3.7.2 software. The clinical and laboratorial results were expressed as medians with interquartile ranges. All RT-qPCR and qPCR assays were performed in experimental duplicates and results were expressed as median ± standard deviations. Significant differences were considered when *p* values were ≤0.05.

## Acknowledgments

The authorsare grateful to AG Costafor aiding in thecytokine network analysis, to Dr AJ Gonçalves for helpin thehistological assays, to Dr H Guedes for the valuable discussions and to the Plataforma de qPCR (RPT09A)/FIOCRUZ.A.M.D.C. holdsa researcher fellowship from CNPqand FAPERJ (CNE).

## Author Contributions

Conceived and designed the experiments: RPR-R, OCM, EFP, AMD-C, AG-S. Performed the experiments: MBP, RPR-R, LR-V, TB-G, MPOO, AFS, LS-C. Analyzed the data: MBP, RPR-R, HGAL, OCM, AMD-C, AG-S. Wrote the paper: AMD-C, AG-S, RRR-P.

---

Table - Multivariate analysis of factors associated with skin lesion size and parasite load in hamsters infected with *Leishmania (Viannia) braziliensis*.

| Variable | Lesion size<br>(mm) |  |  | Parasite load<br>Eq parasites/mg skin |  |  |
| --- | --- | --- | --- | --- | --- | --- |
|  | Correlation | Standard | *p* | Correlation | Standard | *p* |
|  | Coefficient | Error |  | Coefficient | Error |  |
| IgG anti-*Leish* | **.375** | .024 | **.007** | .079 | .059 | .591 |
| Arginase | .343 | .051 | .134 | -.296 | .130 | .234 |
| iNOS | -.061 | .008 | .515 | .037 | .025 | .740 |
| TNF | -.093 | .046 | .596 | -.009 | .177 | .961 |
| IFN-γ | **.576** | .025 | **.017** | .347 | .064 | .172 |
| TGF-β | -.151 | .054 | .197 | -.137 | .133 | .272 |
| IL-10 | -.449 | .035 | .068 | .046 | .091 | .858 |
| IL-6 | .224 | .039 | .252 | **.447** | .097 | **.028** |
| IL-4 | -.082 | .022 | .555 | .261 | .055 | .086 |

\*antileishmanial IgG were evaluated byan ELISA assay; \*\*Parasite load, enzymes and cytokines were evaluated by gene expression (qPCR) according to the Material and methods section. The correlation coefficient values were adjusted considering the inverse CT relationship.

